# Generative Models for Novel Enzymatic Reactions: How Far Do They Go?

**DOI:** 10.64898/2026.11.05.759793

**Authors:** R Ruchir, Srikrishnan Balamuhunthan, Karthik Raman

## Abstract

Enzyme function prediction is a long-standing problem in biology, and deep learning methods have recently become popular to improve performance on this task. While most of these models focus on classifying enzymes or mapping them to known reactions, predicting unknown biochemical reactions remains underexplored. Here we thoroughly examine an approach that generates reactions absent from the training data from the enzyme alone. Existing methods such as BLASTp or CLIPZyme cannot, by design, output reactions that are absent from their training data or were not supplied at inference. The closest prior model, RxnPred, reports high fingerprint-similarity scores on held-out enzymes, a less stringent criterion than exact recovery, but fails to recover the correct answer on unseen reactions. We present a structure-conditioned generative model that outputs a complete reaction, reactants and products, and evaluate it on EnzymeMap_x, a reaction-disjoint benchmark in which no test reaction appears in training, under exact, stereo-respecting scoring. On this benchmark, within 400 sampled candidates, the model generates the complete, correct reaction for 25.0% of the subset of test enzyme–reaction pairs whose reaction is a new combination of metabolites seen in training. Surprisingly, it also correctly generates a few reactions composed of metabolites it has not seen in training (1.6%), though their embeddings are present in the vocabulary. With a lower budget, re-ranking places the correct reaction first for 5.6% of pairs and in the top ten for 12.4%, when the reaction is assumed not to be already known (0.2% and 8.1% without that assumption) along with confidence scores. Assessed more leniently, the model produces at least one correct reaction known for the enzyme (usually memorised from training) for 71.2% of test pairs. Going further, without any retraining, a sequence-only model given the reactants as well as the enzyme produces the correct product set for 9.5% of pairs within five candidates, without re-ranking. We also present head-to-head comparisons on these tasks with BLASTp, DIAMOND, RxnPred and Horizyn. Structure conditioning improves fingerprint similarity on homology-stratified splits, but a matched sequence-only ablation at the best configuration finds no measurable effect on exact recovery. Data coverage and chemical initialisation of the vocabulary seem to matter more than encoder geometry.

## 1 Introduction

Enzymes are the molecular machines that perform biochemical transformations, and understanding their function lets us harness them for the efficient production of industrially relevant compounds and drugs of interest [Sheldon and Woodley, 2018]. Characterising enzyme function has therefore been a longstanding effort in computational biology. A popular expert-curated scheme is the EC (Enzyme Commission) number, a four-level hierarchical classification. Sequencing genes, and hence enzymes, has become very cheap since next-generation sequencing and metagenomics arrived, and NCBI’s Sequence Read Archive alone now holds petabases of raw sequence data [Katz et al., 2022]. Computational biologists have traditionally tackled function prediction by searching massive databases for homologous sequences to approximate the structure and function of a query enzyme. BLASTp and DIAMOND remain powerful tools for this today [Altschul et al., 1990, Buchfink et al., 2021].

Since the deep-learning revolution of the early 2020s, and the availability of predicted structures from tools like AlphaFold2 [Jumper et al., 2021], it has become possible to train neural-network-based models on large corpora of data to help predict enzyme function. Some approaches centred on predicting EC numbers from enzyme sequences include CLEAN, which uses a contrastive-learning approach; CLEAN-Contact, a variant that also uses structural information in the form of contact maps; and HIT-EC, which uses a transformer-based architecture to predict EC numbers hierarchically [Yu et al., 2023, Yang et al., 2024, Dumontet et al., 2026]. However, EC numbers can be difficult to work with: definitions are regularly changed, updated, and added, enzymes do not always fall neatly into these classes, and the distribution across classes is highly uneven. Further, an enzyme assigned a given EC number may only catalyse a subset of the reactions that the label technically covers. So, a more useful, albeit harder, approach is to generate the reaction itself from the sequence. CLIPZyme and Horizyn address the same problem as a screening task: given a reaction, they rank enzymes that could catalyse it, and vice versa [Mikhael et al., 2024, Rocks et al., 2026]. This, however, requires the enzymes or reactions being ranked to already be known at inference time.

Hence, we formulate the problem as a *generative* one, following earlier work from the group that uses an encoder–decoder transformer to generate whole reactions metabolite-by-metabolite given a protein sequence [Balamuhunthan et al., 2026], an approach that is not constrained to any supplied library of reactions. That work reports strong results on a random split: beam search reaches a mean difference-fingerprint similarity of 0.94, with 90% of test entries scored as exact matches (*T* = 1.00) on that metric. On *unseen* reactions, which are the primary application of a generative method, the reported mean structural-fingerprint similarity is 0.79 (median 0.85), and the work states that on unseen reactions the model does not recover the correct answer. Fingerprint similarity also hides detail that exact matching does not: two reactions with a similarity of 1.0 are not guaranteed to be identical, and predictions can differ at exactly the one bond or stereocentre that decides biochemical correctness.

This work builds on that prior work in four ways. We (i) define a reaction-disjoint evaluation taxonomy and exact-match recovery metrics for this class of model; (ii) use more diverse reaction data from EnzymeMap [Heid et al., 2023]; (iii) characterise the architecture more thoroughly, using explicit GNN encoders; and (iv) identify the model’s failure modes and evaluate it more strictly against homology-based methods, retrieval-based methods and the RxnPred baseline itself. The aim is the generative capability that prior work has not yet realised.

## 2 Methods

### 2.1 Data Engineering and Preprocessing

Every enzyme–reaction pair in the test set belongs to one of three difficulty classes that we define for this paper, from easiest to hardest: exact_seen (the reaction, as an ordered multiset of RDKit-canonicalised components, appears in training), recombination (every individual metabolite is seen in training, but not in this combination), or novel_metabolite (at least one metabolite in the reaction is unseen, meaning it appears in no training reaction).

EnzymeMap is a public database of atom-mapped enzymatic reactions [Heid et al., 2023], but most of its entries lack a protein accession. We use EnzymeMap_x as the subset whose enzymes have a UniProt accession [The UniProt Consortium, 2023] with an AlphaFold structure and a sequence embedding, because the original corpus from RxnPred, Rhea [Bansal et al., 2022] + UniProt/SwissProt (Supplementary S7), has sparse reaction coverage. That corpus is used in this paper only for the homology-stratified comparison in §3.1.

We split the pairs 80/10/10. No reaction appears in more than one split, but 818 of the 1,277 test enzymes (64.1%) also occur in training, as each enzyme can have several reactions. The 2,349 test pairs are logically split into 1,275 recombination and 1,074 novel_metabolite pairs, where for 795 of the novel_metabolite pairs the unseen metabolite is a product (novel_product) and in the rest it is a reactant (novel_reactant_only). The vocabulary spans all splits, so the novel metabolites are initialised from their own chemical structure (§2.2.3). Note that this is not open-vocabulary generation, and a reaction-disjoint split does not directly mean the chemistry is new.

76.7% of test pairs (1,801/2,349) could be joined to an EnzymeMap reaction rule, and 99.6% of those use a rule that also occurs among training reactions. The benchmark thus tests recombination of known reaction types over new substrate and co-substrate combinations, not discovery of new reaction types (details in Supplementary S5). 33 test pairs (1.4%, all recombination) are exact reversals of a training reaction; excluding them reduces full-reaction recovery only slightly (Supplementary S12). With stereochemistry ignored, 30 of the 1,072 distinct test reactions (41 pairs, 1.7%) equal a training reaction, so the disjointness is exact only for stereo-aware strings. Table 1 summarises the corpus statistics.

**Table 1:** Corpus statistics for EnzymeMap_x by split. No reaction occurs in more than one split.

| **Quantity** | **Train** | **Val** | **Test** | **Union** |
|---|---|---|---|---|
| Pairs | 18,758 | 2,344 | 2,349 | 23,451 |
| Distinct enzymes | 5,752 | 1,357 | 1,277 | 6,649 |
| Distinct reactions | 8,940 | 1,076 | 1,072 | 11,088 |
| Distinct metabolites | 8,681 | 1,587 | 1,559 | 10,153 |

### 2.2 Model Training

Figure 1 gives an overview of the model described below. We searched hyperparameters with Optuna against validation recall; the search and the resulting final configuration are in Supplementary S3. Our main model was implemented in PyTorch 2.2.1 with PyTorch Geometric 2.7.0 and trained with AdamW, a cosine learning-rate schedule with 20% linear warmup, gradient-norm clipping at 1.0 and bfloat16 autocast, for 12 epochs at batch size 32 (seed 0) on a single NVIDIA A100 GPU. Reaction canonicalisation and scoring used RDKit 2025.09.6 [Landrum, 2025].

**Figure 1:**
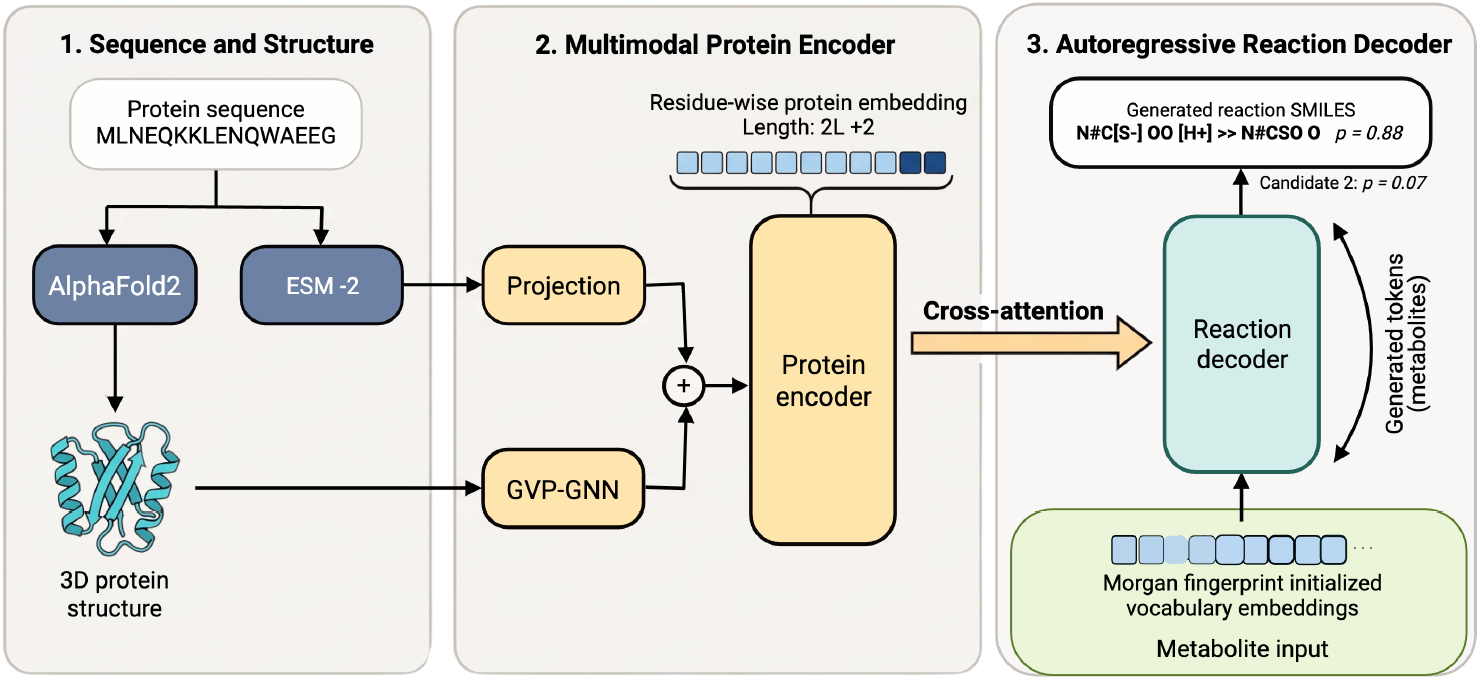
Model overview. ESM-2 [Lin et al., 2023] (sequence) and a GVP-GNN on the AlphaFold2 structure give a residue-wise protein embedding of length 2*L* + 2, which an autoregressive decoder attends to while generating the reaction metabolite by metabolite from Morgan-initialised vocabulary embeddings. SMILES and probabilities are illustrative. Created in BioRender. Raman, K. (2026) https://BioRender.com/pf82na0.

#### 2.2.1 Structural Encoding and Backbone

RxnPred conditions on structure only indirectly, through a frozen ESM-IF1 inverse-folding embedding [Hsu et al., 2022] concatenated onto the ESM-2 sequence stream [Lin et al., 2023]. We instead encode each AlphaFold2 structure as an explicit, trainable *k*-nearest-neighbour graph over C*α*–C*α* distance (*k* = 16) and process it with a graph neural network end to end, alongside the sequence stream. Node and edge features are listed in Supplementary S5.

We selected GVP-GNN [Jing et al., 2021] as the backbone, over a baseline MPNN and a hand-built multi-head geometric-attention architecture (full comparison in Supplementary S1). We stack two GVP-GNN layers, and the network also outputs a max-pooled and a mean-pooled token summarising the whole structure. The GNN’s per-residue features and the ESM-2 per-residue features are concatenated along the token axis, together with the two pooled tokens, giving a sequence of length 2*L* + 2 for an enzyme of length *L*. The decoder reads it through cross-attention.

#### 2.2.2 Additional Structural and Identity-Conditioning Features

The best model also includes pocket-restricted conditioning (P2Rank-derived [Krivák and Hoksza, 2018]) and a separate override for identity reactions. A variant with retrieval augmentation from the training set using Foldseek [van Kempen et al., 2024] was tried and is not part of the model. Full description and ablation are in Supplementary S2.

#### 2.2.3 Chemical Initialisation of Extended-Vocabulary Embeddings

For every token, we standardised an ECFP4 [Rogers and Hahn, 2010] Morgan [Morgan, 1965] fingerprint and a 40-dimensional RDKit descriptor vector, concatenated them, and reduced the result by PCA to *d*_model_ = 512. We then loaded it into the decoder’s token-embedding table as an initialisation that is updated during training, not a frozen one (further details in Supplementary S5). We compared this chemically-informed (“Morgan”) initialisation with ChemBERTa-embedding [Chithrananda et al., 2020] and random initialisations at matched architecture (§3.4).

### 2.3 Inference

We avoid reaction fingerprints as they are not an exact metric and can confuse correct reactions with similar ones, as discussed above.

#### Scoring

Each metabolite is canonicalised with RDKit, and the two sides of a reaction are exactly compared as sorted multisets (so they take stereochemistry into account). *Product-set recovery* compares the product side. *Full-reaction recovery* also requires the generated reactant multiset to match, with reactants and products separated by exactly one ‘>>’.

#### Samples, recovery and recall

The *N* candidates generated for a pair are called samples. *Recovery* is the fraction of test pairs for which any of the first *N* independently sampled candidates matches (best-of-*N* ; duplicates count toward *N* ). *Recall* is the fraction of pairs correctly matched within the top *k* distinct candidates after re-ranking *N* candidates (here we predominantly take *N* = 400; §2.3.2).

#### Uncertainty

Intervals are Wilson 95%, with enzyme-clustered bootstrap intervals that resample the 1,277 enzymes as a check (values in §3.2).

#### 2.3.1 Decoding

Two methods of generating candidates were explored.

*(i) Sampling*.The decoder samples from its trained output distribution after temperature scaling (*T* = 0 is greedy, *T* = 1 the model’s own distribution, *T >* 1 flatter and so favouring rarer tokens). We chose *T* = 1.0 over *T* = 0.7 on the validation split and use it for the test set. Primarily, we sampled 400 candidates (with redundancies) per enzyme–reaction pair at *T* = 1.0 on the test set. As a sensitivity analysis we also sampled at *T* = 1.2 and *T* = 1.5 on the test set, and compared *T* = 1.2 against *T* = 1.0 on validation. Scores were compared across temperatures both unranked (best-of-*N* ) and after re-ranking (§2.3.2).

*(ii) Beam search*. Beam search keeps the *w* highest-likelihood partial outputs at each step and returns all *w*, so the candidate set is small and fixed in size. We decoded widths *w* ∈ { 5, 10, 25, 50} and report both whether the correct reaction appears anywhere in the beam and the top-*k* after re-ranking.

Under sampling, redundancy occurs frequently and grows with budget, for example, at 400 samples the mean number of distinct product sets is only 28.2% of the samples (Supplementary S5). Beam search, by contrast, gives distinct candidates by design. Alternative decode-time diversity strategies (grammar forcing, balancing, resampling) did not outperform the unconstrained control at matched budget (Supplementary S4). The full budget-scaling grid is reported in Supplementary S16 and summarised in §3.2.

#### 2.3.2 Re-ranking

A candidate set lists candidates in sampling order, which carries no information about which is correct. We re-rank each pair’s candidate set using only what is available at inference time: the candidate itself, the candidate set, and the training corpus. We first deduplicate the candidates and then score each distinct candidate on two signals: heavy-atom mass balance between its own reactant and product sides (hydrogen excluded, because proton labelling in the corpus is inconsistent) and how often it was sampled.

Candidates are first filtered by mass balance and then ranked by how often they are sampled. We report two variants. The *assumption-free* variant uses only these two signals. The *novelty-assuming* variant actively searches for new reactions, so it also drops any candidate whose full reaction appears in the training or validation sets. This allows better performance on this benchmark, but a system used for annotation could not assume the answer is new. A learned ranker over the same features did worse than these rules, so we do not use it (Supplementary S13).

#### 2.3.3 Teacher-Forced Product Generation

As a complementary diagnostic, we gave the model the ground-truth reactants at inference so it only had to predict the product from the enzyme. This is forward product prediction, a task that existing models handle well [Schwaller et al., 2019, Sagawa and Kojima, 2025]. It tells us whether the model has learnt chemistry that it can apply once the reactant side is known. It is not the paper’s primary evaluation, which remains the generation of complete reactions from solely the enzyme (§3.4).

### 2.4 Comparison

Aside from RxnPred, other models cannot, by design, predict reactions they have not seen exactly before. So, to approximate a comparison, we compare with three reference points.

*BLASTp and DIAMOND* screen the 1,277 test enzymes against the 5,751-sequence training database, keeping self-hits because the split is enzyme-overlapping, and the product sets of the hit enzymes’ training reactions become the candidate set. Full-reaction recovery is zero for unseen reactions, so only product-set recovery can be compared.

*Horizyn* ranks a supplied library of 11,088 whole reactions that contains the answer, which is an easier task, whereas our model stitches candidates together from a supplied library of metabolites. We compare against Horizyn retrained on EnzymeMap_x and, to provide additional context rather than a fair comparison, against the published checkpoint, whose training data contain 11.0% (118/1,072) of our test reactions.

*RxnPred*. We retrained a sequence-only version on the same dataset (other structural encoders were reported to contribute little). Both models were scored with McNemar’s exact test [McNemar, 1947] and an enzyme-clustered bootstrap interval for the difference. Full protocols are in Supplementary S6 and results are in §3.5 and §3.6.

## 3 Results

All numbers below use the stereo-respecting scoring and the evaluation protocol defined in §2.3, unless a subsection states otherwise. Ranked top-*k* figures are reported in §3.6.

### 3.1 The Effect of Structure

To measure structure’s contribution, we separately trained sequence-only and structure-conditioned models on three homology-stratified splits of the Rhea + UniProt/SwissProt corpus (§2.1), in which test enzymes share at most 30%, 40% or 50% sequence identity with enzymes in training. Structure conditioning improves mean reaction-fingerprint Tanimoto similarity at every threshold tested: 0.689 → 0.733 (+0.044) at 30% identity, 0.872 → 0.897 (+0.025) at 40%, and 0.926 → 0.961 (+0.035) at 50%, with both models greedy-decoded.

We then tested the structure ablation directly, training two models that differed only in whether the structural channel was included as input (same hyperparameters, Morgan initialisation, seed and 12 epochs). Recovery within 400 samples was indistinguishable for both full-reaction and product-set recovery (McNemar *p* = 0.75 and *p* = 0.34; Supplementary S11), with opposite signs on the two metrics. Retraining the structure model recovered 316 test pairs (13.45%) in full instead of the best checkpoint’s 336 (14.30%). That is as large as the product-set difference ( −0.85) and three times the full-reaction difference (+0.26). We therefore read the fingerprint-similarity gains above as real but small, and treat structure’s contribution to exact recovery as not established rather than excluded. In short, structure improves similarity but not exact recovery.

### 3.2 Recovery on Reaction-Disjoint Test Pairs

Each of the 2,349 pairs is either a recombination of seen metabolites or involves an unseen one. Prior generative work on this task reported novel (recombinant) reactions being emitted, but at low frequency [Balamuhunthan et al., 2026]. With only the enzyme supplied, the model recovers the complete reaction for 14.30% of pairs within 400 samples (336/2,349) and the exact product set for 27.93% (Figure 2; Supplementary S16). We apply no re-ranking to any number in this section, so these numbers show the model’s raw predictive power and are affected by redundant generation (duplicate candidates). About half of the product-set recoveries also reproduce the reactants (336 of 656), and the reactant side alone is recovered for 29.0% of pairs (682). Recovery is still rising at 400 samples (Figure 2).

**Figure 2:**
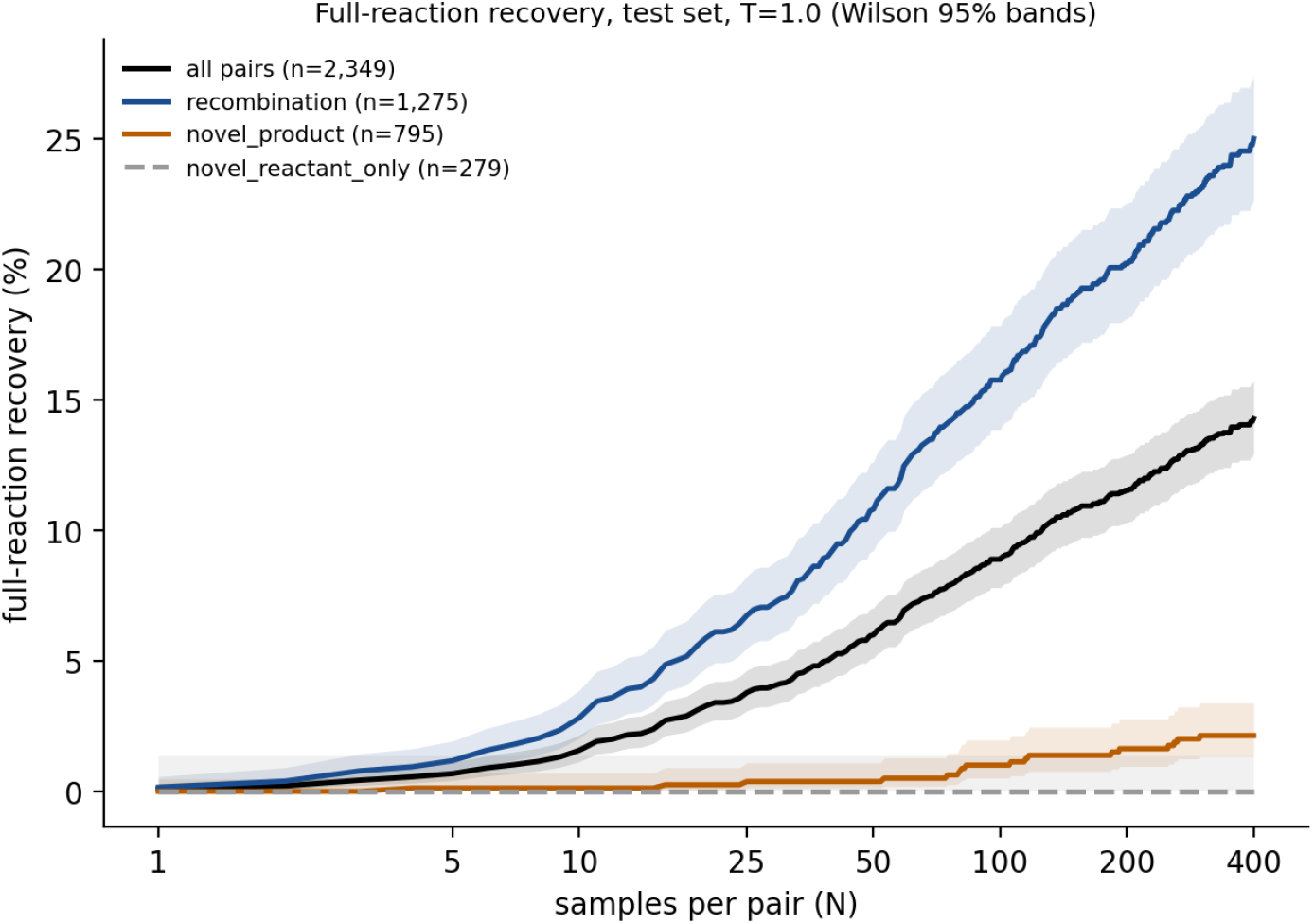
Full-reaction recovery within *N* samples on the test set (*n* = 2,349, *T* = 1.0) for all pairs, recombination pairs and the two novel-metabolite subclasses. Wilson 95% bands shown; values at selected *N* are in Supplementary S16.

Enzyme-clustered bootstrap intervals are only slightly wider than the Wilson intervals (Supplementary S12). Independent sampling runs at identical settings differ by up to 1.0 percentage points for product-set recovery and 0.3 for full-reaction recovery.

#### Sampling temperature and beam search

Temperature sets how far the decoder strays from its most likely tokens: higher values flatten the output distribution, so rarer tokens are sampled more often. We found that higher temperature helps generate more novel reactions. Full-reaction recovery peaks at *T* = 1.2 and falls at *T* = 1.5, where more of the candidates are invalid reactions. Recovery of reactions with novel metabolites falls slightly but consistently as temperature rises. After ranking, *T* = 1.2 beats *T* = 1.0 from about *k* = 25 onward, while *T* = 1.0 is best for the first few ranked candidates. Therefore, modestly higher temperatures increase the variance, and with it the number of unique reactions emitted, but if the temperature is too high the reactions become malformed and invalid. Beam search returns only the highest-likelihood outputs. At every width tested it rarely contained the correct full reaction, and for product prediction it recovered less than sampling at 25 candidates. We therefore use sampling throughout (both tables are in Supplementary S4).

### 3.3 Recombination versus Novel-Metabolite Generalisation

Plotting performance against sampling budget for each of the difficulty classes in §2.1 (`recombination`, `novel_metabolite`, and its subclasses `novel_product` and `novel_reactant_only`) shows that recovery differs dramatically between them (Figure 2; full table in Supplementary S16).

Within 400 samples, a quarter of recombination pairs are correctly predicted (319 of 1,275; 25.02%). Pairs that need an unseen product metabolite are recovered much more rarely (17 of 795, 2.14%). None of the 279 reactions whose unseen metabolite is a reactant is generated, so the model cannot reliably predict a reaction containing an unseen reactant. The gap between classes widens with the sampling budget. That the novel-metabolite rate is non-trivial at all, given that these metabolites receive no positive training signal, is examined in §3.6.

### 3.4 Vocabulary Initialisation and Teacher-Forced Product Generation

#### Vocabulary initialisation

We compared three initialisations at matched architecture (§2.2.3). Product-set recovery within 25 samples is 8.39% with the Morgan-fingerprint initialisation, 6.90% with ChemBERTa embeddings and 6.13% with random initialisation. In paired McNemar tests, Morgan beats random (*p* = 0.0002) and ChemBERTa (*p* = 0.026), while they do not differ significantly from each other (*p* = 0.234). A possible reason is that token-level language-model embeddings do not carry the stoichiometric information, which is essential for generating balanced chemical reactions.

#### Teacher-forced product generation

In this diagnostic we supply the ground-truth reactants and ‘>>’ (§2.3.3) and the model predicts only the product. On the sequence-only model (beam width 5, 2,349 test pairs), top-1 product-set recovery rises from 0.72% (17 pairs) when generating freely to 2.94% (69) with teacher forcing, and best-of-5 recovery rises from 2.34% (55) to 9.54% (224). This shows that the model can complete a reaction when part of the answer is given, indicating that it may be capable of completing reactions. These are product-set numbers only, and full-reaction generation remains the primary evaluation.

### 3.5 Generalisation

#### Does recovery read the enzyme?

To ensure that the model is gleaning the reaction from the enzyme input and not reaction frequency or sequence homology, we:

##### Shuffled enzymes

We gave each test pair a different enzyme (a derangement of the enzyme assignments, with the same checkpoint and scorer). Recovery within 400 samples fell to 0.77% for product sets and 0.38% for full reactions; a baseline with no enzyme and no model, which just ranks training product sets by frequency, reaches 0.81%. Of the pairs recovered under either condition, 328 were recovered only with the true enzyme and 1 only with the shuffled one (exact McNemar *p* = 6 ×10^*−97*^).

##### Homology

If the model were just copying close relatives, recovery would rise with similarity to the training set, but no such trend exists. Across bins of BLAST identity to the nearest training enzyme, recovery ranges from 10.7% to 17.3% with no monotone trend, and the highest-identity bin ( ≥90%) is among the lowest (Supplementary S12). Of the recovered product sets, 83% never occur anywhere in training, so the model is not simply reproducing a memorised answer.

#### Scoring at Enzyme Level

An enzyme can catalyse more than one reaction, so we argue that a candidate matching *any* known reaction of the enzyme, not only the pair’s own target, should count as a correct prediction. We therefore extended the ground-truth hits to any reaction that the enzyme catalyses in train, validation or test. Under this rule, test recovery rises from 14.3% (336 of 2,349) to 71.2% (1,673 of 2,349). Almost all of that rise is memorisation: 98.5% of the newly credited pairs are credited only because the *same* enzyme also has a memorised training reaction.

##### Multi-reaction enzymes

Of the 1,277 test enzymes, 586 have two or more known reactions. At 400 samples, all of their reactions are recovered for only 47 of these enzymes (8.0%), although at least one reaction is recovered for about 21% of test enzymes. On average 13.2% of a multi-reaction test enzyme’s reactions are recovered, close to the overall test rate of 14.3%, so the low rate of recovering all reactions may largely reflect the need for every reaction to be hit (Supplementary S8).

We report this only as a check on our own protocol, not as an alternative way to measure recovery. For the 818 of 1,277 test enzymes that also occur in training (§2.1), any enzyme-level rule mostly reflects what the model memorised about the enzyme. Every other number in this paper is therefore scored against the pair’s own target only (Supplementary S8).

### 3.6 Re-Ranking Generated Candidates

Among the 336 recovered pairs, the median first hit is sample 61 (quartiles 24 and 147), and only 37 are hit within the first ten samples. The mean reciprocal rank over all pairs is 0.0066. Recovery in sampling order (Figure 2) therefore depends on the order of the candidate set as well as on what it contains.

#### Benefit of re-ranking

We ranked each 400-sample candidate set with the procedure of §2.3.2. Under the novelty assumption, the correct ground-truth reaction comes first for 5.62% of the 2,349 test pairs and appears in the top ten for 12.39% (291 pairs). The originally sampled order is far lower at every *k* (Figure 3). Ten ranked candidates therefore hold 291 of the 336 reactions that the whole candidate set contains (86.6%). Mass balance, sampling frequency and the novelty filter only work in combination. By *k* = 50 re-ranking reaches 333 of the 336, so ordering stops being the limit.

**Figure 3:**
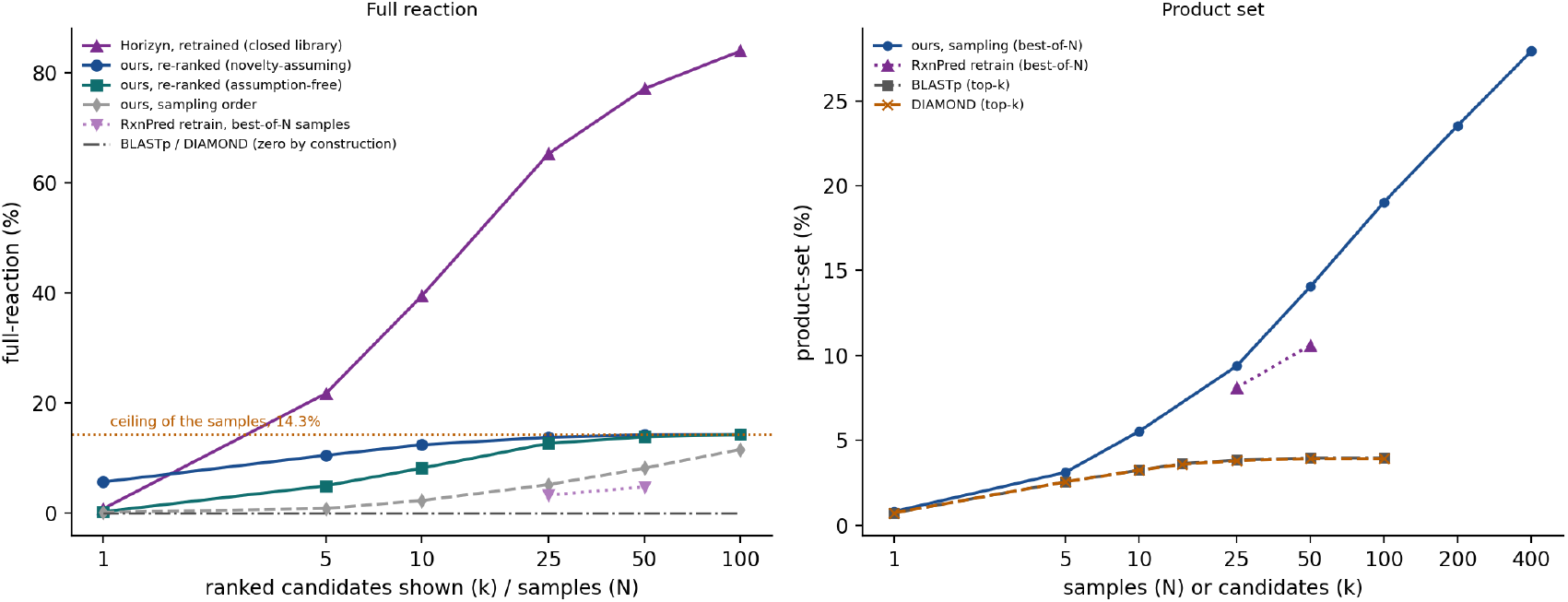
Comparison with other methods on the test set (*n* = 2,349). *Left:* full-reaction recall when the top *k* ranked candidates are shown. Our generator is shown in random sampling order, re-ranked under the novelty assumption, and re-ranked without it (assumption-free). Horizyn ranks a closed library from train, validation and test that contains the answer. BLASTp and DIAMOND only return training reactions, so their full-reaction recovery on our test set is zero by construction. The RxnPred retrain is shown at 25 and 50 unranked samples. *Right:* product-set recovery, the only level at which homology search can be compared. Ours and RxnPred are best-of-*N* over samples, while BLASTp and DIAMOND are recall over their top *k* candidates.

#### Benchmark measurement versus exploratory use

The novelty filter removes any reactions of that enzyme that were seen during training or validation. It accounts for most of the high top-1 performance, and without it top-1 is 5 pairs (0.21%) instead of 132. The assumption-free variant naturally emits many reactions already seen in training, since the data is not enzyme-disjoint. It reaches a top-10 of 191 pairs (8.13%), 3.6 times the sampling-order top-10, and the two variants converge by *k* = 100.

Of the 132 top-1 recoveries, 129 are recombination pairs (out of 1,275) and 3 are novel-metabolite pairs (out of 1,074). At top-10 the split is 275 and 16, which is 21.6% of recombination pairs. Product-set recall under the same ranking is in Supplementary S9.

#### Comparison with external methods

##### Homology search

We screened the 1,277 test enzymes against the 5,751 training sequences with BLASTp and DIAMOND (protocol in §2.4) and used the product sets of the best hits’ training reactions as candidates. Only product sets are comparable, since full-reaction recovery is zero on this test split. Both curves plateau near 4%, against 9.37% for our model at 25 samples, so our product-set recovery is always higher (Figure 3).

##### RxnPred retrain

We retrained RxnPred’s architecture on the same data (protocol in §2.4). Our model recovers more product sets at both 25 and 50 samples (*p* = 0.0016 and *p* = 0.0006; Supplementary S15). For full reactions the difference is not significant at 25 samples (*p* = 0.14) and is significant at 50 (*p* = 0.043).

The models differ in multiple ways, including initialisation, structure conditioning and hyperparameters.

##### Horizyn

Horizyn ranks a limited library of 11,088 reactions that is given to contain the answer, in contrast to our generator, which ranks across an unlimited reaction space. We compared recall at rank *k* against Horizyn retrained on our data (Supplementary S15). With the novelty assumption our recall is ahead at *k* = 1 (5.62% against 0.77%) and behind at every larger *k* (for example 12.39% against 39.46% at *k* = 10). Without it we are behind at every *k*, including *k* = 1 (0.21% against 0.77%). The published Horizyn checkpoint, trained on far more data and on 11.0% of our test reactions (118 of 1,072), reaches 18.35% and 46.70% at *k* = 1 and *k* = 5, so we make no claim against it.

The two models work in different settings, so this contrast does not rank them. A retrieval model can only return reactions in its library, and within that regime it is far stronger at recall. The generator’s top-ranked candidate is correct for 5.6% of pairs when novel chemistry is assumed, in a setting where no library is supplied and the answer is, by construction, a reaction absent from training. Pooling the candidates of several checkpoints (an ensemble) enlarges the candidate set but does not improve ranking at small *k* over a single model; the analysis is in Supplementary S10.

#### Successful examples

Figure 4 shows three recovered reactions, and Supplementary S14 shows three more. One is the prenylation of a cyclic dipeptide by dimethylallyl diphosphate, whose product is absent from training. The other two are recombination reactions: a transamination of glyoxylate by glutamate and the decarboxylation of arginine to agmatine. All three are complete, stereo-specified biochemistry. Each first appears between sample 25 and sample 132 of its candidate set, so a reader who inspects only the first few samples of the unranked output would see none of them. The other three have products absent from training: the glucosylation of an isoflavone by UDP-glucose, the glucuronidation of an acetylated tryptamine by UDP-glucuronate, and the oxidation of a polyhydroxylated steroid by molecular oxygen (Supplementary S14).

**Figure 4:**
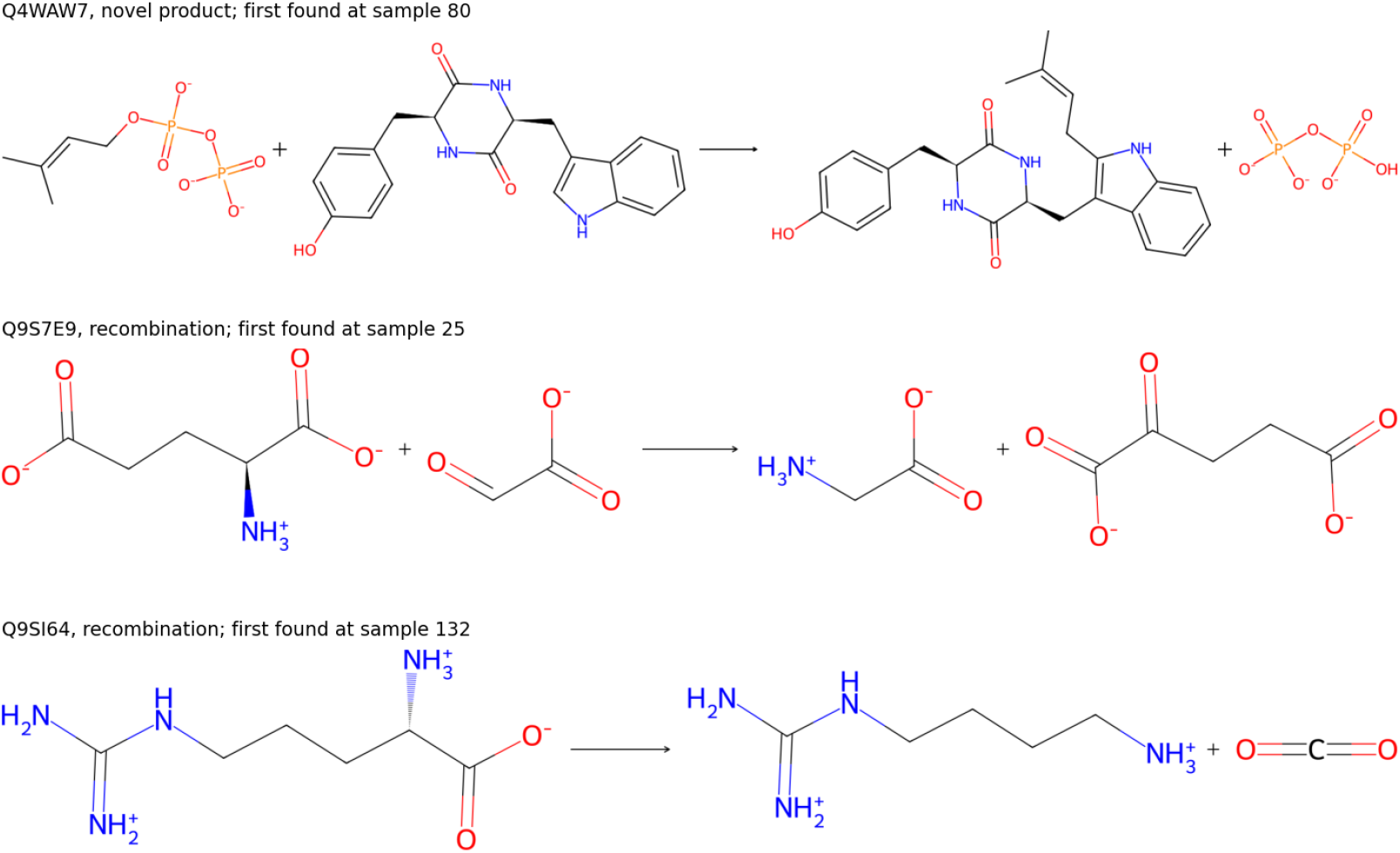
Recovered full reactions from the test set, with the UniProt entry, the difficulty class and the sample at which the reaction first appeared among the 400 candidates.

Our hypothesis, untested here, is that such recoveries are templated group transfers or oxidations whose donor or co-substrate half is dense in training (dimethylallyl diphosphate, UDP-sugars, glutamate and glyoxylate, O_2_), so the model need only place one new acceptor.

#### Metabolites absent from training

The novel_product recovery reported in §3.3 is surprising. These pairs require the decoder to emit a product metabolite that was never a training target. We now discuss two relevant observations.

*Observation 1*. Of the 795 test pairs whose product includes a metabolite absent from training, the model recovered the full reaction for 17 (2.1%) within 400 samples and the product set for 43 (5.4%). All 17 involve a product that is not in training even after stereochemistry and charge are ignored. Also interesting is that none of the 117 pairs whose unseen product is only a stereo or protonation variant of a seen metabolite is correctly predicted, so the successful cases need a sufficiently distinct molecular structure.

*Observation 2*. The embedding rows of never-trained tokens stay close to their initialisation (cosine similarity 0.9959, against 0.9988 for trained rows). Technically, since the output projection is tied to the embedding table, these rows still receive negative-class gradient at every training step, but they are never trained toward any output. The vocabulary holds these metabolites with structure-derived embeddings at inference (§2.1), so the model selects them from that vocabulary. Structure-derived initialisation may therefore contribute to recovering unobserved metabolites, though here we do not isolate the mechanism.

## 4 Discussion

Incorporating fine-grained structural information improved fingerprint similarity modestly, but a matched ablation finds no measurable effect on exact recovery. One possible reason is scale of supervision, as a graph encoder trained on a few thousand enzyme–reaction pairs has far less information to learn geometric detail from than the sequence stream, for which ESM-2 [Lin et al., 2023] was pretrained on a much larger corpus. The improved performance reported in §3.2–3.3 came mostly from extending the vocabulary and informing its embeddings chemically (§2.1, §2.2.3) rather than from the encoder. Here, data coverage and representation mattered more than fine-grained geometry.

The difference in performance between recombination and novel-metabolite recovery (§3.3) fits how the model is supervised. Each training example teaches the model to choose amongst metabolites it has seen as positive targets, so recombining them into new reactions is highly achievable, but building a reaction around a metabolite that was never a positive target is far more difficult. The embedding observation in §3.6 implies that the chemically informed initialisation may carry enough structure for the model to occasionally place such a metabolite correctly.

When searching for the best candidates through the sampling results, we use a simple intelligent reranker which has two variants, where the assumption-free variant tries to annotate the enzyme, while the other looks for plausible unseen reactions it could catalyse. On comparing with existing methods, retrieval models perform annotation much better than we do (Horizyn), albeit within a given pool of candidates known at the time of inference. This benchmark and generative framework attempts to find new possible reactions that known enzymes could catalyse, and we show that it frequently succeeds in predicting new reactions. A longer form of these arguments is in Supplementary S17.

Taken together, these results describe a model whose chemical generalisation is largely compositional rather than extrapolative: it is substantially better at recombining chemistry it has seen in new ways than at reasoning about chemistry it has not seen before. It is unclear whether the limits of performance are due to the architecture or to the training information available to it.

### 4.1 Future Work

Attention maps and other explainability analyses could show which parts of the enzyme the model attends to and guide improvements. Testing on established benchmarks, including a leave-one-organism-out split and EC-based or rule-based splits, would separate enzyme-level from reaction-type generalisation and inform the applicability of the approach. The model should also be conditioned on pH, temperature, and cofactor and metal-ion availability, which it currently ignores. Also, balance-aware losses or masked decoding, along with other approaches to increase diversity, could help increase the variety of reactions produced per sample.

### 4.2 Limitations

The vocabulary is transductive: test-only metabolites have embedding rows initialised from their own chemical structure before training (§2.1, §2.2.3), so novel-metabolite recovery is selection from a supplied vocabulary, not truly open-vocabulary generation. This can be achieved using alternate tokenizers, however. Since many of the test enzymes occur in training, enzyme-level generalisation is not cleanly characterised, and an enzyme-level scoring rule would inflate recovery from 14.3% to 71.2% (§3.5). The reverse of the true reaction appears among the candidates for 8.1% of test pairs, but most of those are also forward hits, and these are reported separately. In addition, the benchmark does not split by known chemistry, where the majority of test reactions are governed by a rule already present in training (§2.1), and recovery is far higher for recombination than for novel-metabolite pairs (§3.3), so the reported numbers characterise recombination of known reaction types more than discovery of new ones.

Further, the ranking results are heavily supported by the novelty assumption (§3.6), and the mechanism behind novel-metabolite recovery is not isolated. As expected, the generator learned mass balance only as a soft preference (22.9% of distinct candidates but 45.3% of samples are balanced, Supplementary S9), and the majority of test enzymes do not have the correct candidate among the 400 samples (§3.2). Recovery is therefore limited by raw generative capacity as well as by re-ranking strategies. Overall recovery (14.3% full-reaction, 27.9% product-set, best-of-400) is well above the homology and frequency baselines but is still short of a reliably usable rate.

### 4.3 Conclusion

We trained a structure-conditioned generative model to predict possible reactions from the enzyme alone and evaluated it on a reaction-disjoint benchmark under exact scoring. It recovered the complete reaction for 25.0% of recombination reactions within 400 samples. Controls indicate that these predictions depend on the enzyme and are not products of homology or reaction frequency. Notably, reactions that need a metabolite absent from training are recovered far less often (1.6%) but still above zero. Re-ranking the candidate set without ground-truth reactants places the correct full reaction first for 5.6% of pairs when the answer is assumed to be a never-before-seen reaction. A matched ablation finds no measurable effect of structure on exact recovery. This work primarily serves to demonstrate the potential of this transformer-based approach in the annotation of enzymes and, more importantly, the discovery of novel reactions they can catalyse.

## Acknowledgements

We acknowledge the National Supercomputing Mission (NSM) for providing computing resources of ‘PARAM RUDRA’ at the P G Senapathy Center for Computer Resources, Indian Institute of Technology Madras, Chennai, Tamil Nadu 600036, which is implemented by C-DAC and supported by the Ministry of Electronics and Information Technology (MeitY) and the Department of Science and Technology (DST), Government of India. The authors used AI-based language editing tools (ChatGPT 4.0 and Grammarly) to improve the grammar and readability of the manuscript. No AI tools were used for data analysis, interpretation, or drawing scientific conclusions.

## Data Availability

Scripts for training and evaluation are available at https://github.com/Ruchir-r/GenEnzyme, along with the datasets and sampled candidate pools.

## Notes

### Competing Interest Statement

The authors have declared no competing interest.

