## Supplementary Material for "Generative Models for Novel Enzymatic Reactions: How Far Do They Go?"

S1–S6 give details of the model, decoding and comparisons, and S7 describes the corpora. S8–S12, S15 and S16 hold the tables and checks behind the reported numbers, S13 lists negative results, S14 shows recovered examples, and S17 expands the Discussion. Section numbers such as §3.1 refer to the main paper.

#### S1. GNN Backbone Selection

Three backbones were compared over the structural graph of main paper §2.2.1: MPNN, a hand-built multi-head geometric-attention architecture, and GVP-GNN [Jing et al., 2021]. GVP-GNN gave the best mean Tanimoto (0.9303, against 0.9262 for the hand-built geometric-attention architecture, 0.9207 for MPNN and 0.9123 sequence-only) and is used throughout the main text. These are single-run values with no variance estimate.

This comparison was run on the Rhea [Bansal et al., 2022] + UniProt/SwissProt [The UniProt Consortium, 2023] test split ($n$ = 23,403, S7), which is enzyme-disjoint but *not* reaction-disjoint: 98.6% of its reactions also occur in training. So it compares backbones mostly on seen chemistry, and the result should not be read as reaction-disjoint. The homology-stratified splits in main paper §3.1 test the same question under tighter control.

#### S2. Auxiliary Structural and Identity-Conditioning Features

Two additions were made to the GVP-GNN backbone one after the other, and each was kept only if it improved on the one before: pocket-restricted conditioning (an additive bias and a pooling token built from P2Rank [Krivák and Hoksza, 2018] pocket probabilities; a multiplicative gate on the pocket signal trained to a negative weight and was dropped), and an identity-aware override that lets the model detect near-identical training enzymes and copy reactants to products when triggered. Focal cross-entropy loss ($\gamma = 2$) was the largest single gain. On the Rhea + UniProt/SwissProt test split mean Tanimoto rises from 0.9303 (GVP alone) to 0.9322 with pocket conditioning and 0.9361 with the identity-aware override and focal loss. A third addition, Foldseek-based retrieval augmentation (concatenating the reactions of the five most structurally similar training proteins onto the encoder output), was tried as a separate experiment, gave +0.007 mean Tanimoto, and is not part of the deployed model.

The identity override's loss weight is 0 in the best configuration (S3), so it is switched off for every free-running result in the main paper (that is, generation of the full reaction from the enzyme alone, with no reactants supplied); it only influenced the conditioning features chosen during the search.

#### S3. Hyperparameter Search and Final Configuration

Hyperparameters were searched with Optuna (TPE sampler). An initial pass that minimised validation loss ranked the initialisation options wrongly: ChemBERTa reached lower loss but fewer product hits than Morgan [Morgan, 1965, Rogers and Hahn, 2010] initialisation at matched budget. A second pass searched directly against validation recall and is the source of the best configuration, so the main text describes the search as targeting recall.

That configuration, `cbv2_t010`: Morgan initialisation, $d_{\text{model}} = 512$, 8 heads, 4 decoder layers, feed-forward 1536, dropout 0.2, two GVP layers, base learning rate $3.73 \times 10^{-4}$, GNN learning rate $1.72 \times 10^{-3}$, weight decay $1.76 \times 10^{-3}$, cosine schedule with 20% warm-up, batch size 32, focal-loss $\gamma = 2$, identity-loss weight 0. Random initialisation reached 3.29% validation recall against Morgan's 11.05%, with no overlap between trials, and was dropped from later search spaces.

**Software and hardware.** Models were trained with a custom loop in Python 3.10.13 under Slurm 23.11.10 on a RHEL-family 8 cluster (kernel 4.18) with CUDA 12.1 and cuDNN 8.9.

### S4. Decoding Strategies

Beyond temperature-scaled sampling (main paper §2.3.1), three decode-time diversity strategies were tried: grammar forcing, reachability-gated balancing, and resampling of already-seen candidates. None outperformed unconstrained sampling at matched budget, and resampling also failed to complete at scale. None is used in the best model.

**Temperature on validation.** The temperature used for free-running full-reaction recovery was chosen on the validation split ($T = 1.0$ over $T = 0.7$). A later validation sweep compared $T = 1.2$ with $T = 1.0$: product-set best-of-400 recovery was 32.34% against 29.61%, while the first-sample hit rate was lower at $T = 1.2$ (0.68% against 1.11%). $T = 1.5$ could not be run on validation and is a test-only sensitivity analysis (main paper §3.2). The test-set temperature table is in S5.

**Beam search.** Beam search returns the few highest-likelihood outputs instead of sampling, and is the obvious choice when only the top candidate will be inspected. It rarely contains the correct full reaction at any width tested.

Table 1: Beam search, test $n$ = 2,349. "In beam" counts pairs whose correct full reaction appears anywhere in the beam; "malformed" is the share of beam outputs containing no `>>`. For comparison, 400 samples recover 336 pairs, with re-ranked top-1 of 5 (assumption-free) and 132 (novelty-assuming).

| width | correct in beam | malformed outputs | top-1 after re-ranking: assumption-free | top-1 after re-ranking: novelty-assuming |
|---|---|---|---|---|
| 5 | 25 | 10.9% | 2 | 24 |
| 10 | 33 | 14.4% | 7 | 31 |
| 25 | 32 | 18.4% | 6 | 25 |
| 50 | 28 | 20.7% | 8 | 21 |

In exact beam order the correct reaction ranks first for at most one pair at any width. Widths above 50 were not scored. On the product side beam search is also behind sampling at a matched 25 candidates.

Table 2: Product-set recovery from 25 candidates, test $n$ = 2,349. The two sampling rows are independent runs at $T = 1.0$.

| 25 candidates from | product-set recovery |
|---|---|
| beam search, width 25 | 7.02% |
| sampling, run 1 (S16) | 9.37% |
| sampling, run 2 (S12) | 10.34% |

Because beam search loses on both metrics we use sampling throughout. It does, however, return distinct candidates by design, where sampling does not (S5).

### S5. Model, Decoding and Benchmark Specifications

Details that the main text mentions but does not describe.

**Structural encoder features.** Node scalar features are the AlphaFold2 per-residue confidence score (pLDDT), backbone torsion angles, and radial-basis-function encodings of distances to the backbone N, C, O and the side-chain centroid. Node vector features are the corresponding unit direction vectors. We add one enzyme-specific feature, a ligand-binding-pocket residue indicator from P2Rank [Krivák and Hoksza, 2018]. Edges carry a radial-basis-function distance encoding and a unit direction vector. GVP-GNN keeps separate scalar and geometric vector channels per residue and restricts every operation on the vector channel to linear combination and an $L_2$ norm, which is sufficient to make the vector output equivariant, and the scalar output invariant, under any rotation or reflection of the input coordinates [Jing et al., 2021].

**Chemical initialisation.** The 40-dimensional RDKit descriptor vector comprises atom counts, molecular weight, charge, ring counts, H-bond donor and acceptor counts, TPSA, MolLogP, and related descriptors. Special tokens keep random initialisation. The output projection shares its weight matrix with the token-embedding table.

**Decoding.** Under sampling, the mean number of distinct product sets is 64.4% of 25 samples, 53.9% of 50, and 28.2% of 400, so redundancy grows with budget. We used bfloat16 autocast at every decoding budget; one small-scale fp32 check found no meaningful difference.

**Benchmark overlap.** Of the test pairs joined to an EnzymeMap reaction rule (1,801), 1,793 (99.6%) use a rule that also occurs among training reactions (253 rules in train, 93 in test; EC class to the third level seen for 1,800/1,801). 8.5% of test pairs (199) have a product set that occurs on the product side of some training reaction, and 17.4% a reactant set. This supports the main text's statement that the benchmark tests recombination of known reaction types rather than discovery of new ones.

**Sampling temperature.** Table 3 gives the test-set effect of sampling temperature, summarised in main paper §3.2. Raising $T$ increases unranked product-set recovery but lowers precision after re-ranking.

Table 3: Effect of sampling temperature, test $n = 2{,}349$, 400 samples per pair at each $T$. Best-of-400 columns give unranked recovery; the remaining columns give the number of pairs whose full reaction is in the top $k$ after re-ranking, novelty-assuming and assumption-free.

| | **Best-of-400** | | **Novelty-assuming** | | | **Assumption-free** | | |
|---|---|---|---|---|---|---|---|---|
| $T$ | product-set | full | $k=1$ | $k=5$ | $k=10$ | $k=1$ | $k=5$ | $k=10$ |
| 1.0 | 27.9% | 336 (14.3%) | 132 | 246 | 291 | 5 | 116 | 191 |
| 1.2 | 30.9% | 345 (14.7%) | 120 | 234 | 284 | 4 | 114 | 189 |
| 1.5 | 31.4% | 298 (12.7%) | 99 | 192 | 234 | 5 | 93 | 163 |

### S6. Comparison Protocols

The three reference points of main paper §2.4, with the settings that the main text leaves out. Results for all three are in main paper §3.6, with the tables in S15.

*BLASTp and DIAMOND.* We screened the 1,277 test enzymes against the 5,751-sequence training database in reverse, with BLASTp (`-evalue 10 -max_target_seqs 500`) and DIAMOND (`--very-sensitive`), keeping self-hits because the split is enzyme-overlapping: 818 of the 1,277 test enzymes also occur in training. For each query enzyme we take as the candidate set the product sets of the hit enzymes' training reactions, best hit first. Because no test reaction occurs in training, full-reaction recovery is zero, so only product-set recovery can be compared.

*Horizyn.* Horizyn ranks a supplied library of 11,088 whole reactions that contains the answer, so its recall is over complete reactions and the like-for-like quantity for our model is full-reaction recall after re-ranking. Its task is also easier than ours, since it ranks a closed library while we sample openly and our samples include duplicates. We compare against Horizyn retrained on EnzymeMap_x and against the published checkpoint, at ranks $k \in \{1, 5, 10, 25, 50, 100\}$, with and without the novelty assumption. Because 11.0% (118/1,072) of the test reactions are in the published checkpoint's training data, the second comparison is only for context and is not a controlled comparison.

*RxnPred.* We retrained RxnPred's architecture, sequence-only, on EnzymeMap_x in one run with no tuning. The retrain has 41.2M parameters, ESM-2 [Lin et al., 2023] 480-d input, an extended vocabulary and trainable ChemBERTa-initialised embeddings, and was trained for 25 epochs. We scored both models on the same 2,349 pairs, using McNemar's exact test [McNemar, 1947] and an enzyme-clustered bootstrap interval for the difference. The two models differ in initialisation, structure conditioning and hyperparameters, so the comparison does not isolate structure.

### S7. Corpora

EnzymeMap_x, defined in main paper §2.1, supplies every number in this paper except the homology-stratified comparison in main paper §3.1 and the backbone and feature selection in S1 and S2, which use the Rhea + UniProt/SwissProt corpus described here. That corpus is the one used by RxnPred; we stopped using it because its reaction coverage is sparse.

We downloaded protein sequences, with their EC numbers and Rhea annotations [Bansal et al., 2022], from SwissProt [The UniProt Consortium, 2023] on 15 September 2025. We removed sequences shorter than 30 or longer than 1,024 amino acids and kept only entries with between one and ten Rhea associations, following RxnPred [Balamuhunthan et al., 2026]. We used only the forward (left-to-right) version of each reaction, filtered out reactions without a valid Rhea SMILES string, and canonicalised each metabolite's SMILES string with RDKit [Landrum, 2025]. We restricted AlphaFold2 structures to the AFDB SwissProt-structures subset [Varadi et al., 2022]. The splits are homology-stratified by MMseqs2 clustering [Steinegger and Söding, 2017] at 30%, 40% and 50% sequence identity, so results on this corpus are enzyme-disjoint but not reaction-disjoint.

### S8. Enzyme-Level Recovery Under an Any-Known-Reaction Rule

Supporting detail for the enzyme-overlap check in main paper §3.5. A candidate is scored correct here if its canonical full reaction equals *any* reaction known for that enzyme in any split, using an enzyme→known-reactions lookup spanning train, validation and test and built separately from the pair-level ground truth used everywhere else in this paper. Inference was extended to the 5,372 enzymes outside the test set (6,649 in the corpus minus the 1,277 test enzymes), which had no candidate set because the main runs covered test pairs only (400 samples, $T = 1.0$); the 1,277 test enzymes reused their existing candidate sets. About 30 of the newly sampled enzymes (under 1%) logged a missing structure and were not stratified separately.

**Known reactions present in the candidate set, by the known reaction's own split** (23,451 known enzyme–reaction pairs; % of pairs for which that pair's reaction appears in the enzyme's candidate set):

| samples | train (18,758) | val (2,344) | test (2,349) | all (23,451) |
|---|---|---|---|---|
| 1 | 6.63% | 0.13% | 0.17% | 5.33% |
| 25 | 46.20% | 4.01% (94) | 3.87% (91) | 37.75% |
| 100 | 64.78% | 7.76% (182) | 8.81% (207) | 53.48% |
| 400 | 77.57% (14,551) | 13.44% (315) | 14.30% (336) | 64.82% (15,202) |

The test figure at 400 samples (336/2,349) reproduces the independent pair-level audit behind the main paper's free-running recovery exactly. The two were computed from different code paths, so this only checks that the scoring is consistent and is not new evidence.

**Enzyme-level recovery at 400 samples.** A multi-reaction enzyme has two or more known reactions in that split. The "≥1 recovered" column covers all enzymes in the split; "all recovered" counts multi-reaction enzymes with every one of their reactions recovered.

| split | enzymes | multi-reaction | ≥1 recovered | all recovered | mean fraction recovered |
|---|---|---|---|---|---|
| train | 5,752 | 3,979 | 92.37% | 64.94% (2,584) | 84.70% |
| val | 1,357 | 643 | 20.78% | 2.02% (13) | 11.07% |
| test | 1,277 | 586 | 21.06% | 8.02% (47) | 13.21% |
| all | 6,649 | 4,751 | 83.20% | 47.84% (2,273) | 70.82% |

The overall figures are driven by training reactions, which the model memorised; only the validation and test rows measure generalisation. On test, the mean fraction recovered per multi-reaction enzyme (13.21%) and the pooled rate over all pairs (14.30%) weight enzymes differently and are only loosely comparable. Pooled over pairs, reactions of single-reaction test enzymes are recovered more often (153 of 691 pairs, 22.1%) than those of multi-reaction test enzymes (183 of 1,658 pairs, 11.0%), so the low "all recovered" rate may reflect the need to hit every reaction of the enzyme.

**Two controls on what the rule measures.** Scoring a candidate set against a *different* enzyme's known reactions gives 0.61% (train), 0.20% (val) and 0.21% (test), so the candidate sets are enzyme-specific. Consistent with that, two test pairs sharing an enzyme draw much more similar candidate sets (mean Jaccard 0.199) than test pairs of different enzymes (0.001). Of the 1,337 test pairs newly credited by the rule, 1,317 are credited through the enzyme's own memorised training reaction, 166 through another test reaction of the enzyme and 76 through a validation reaction (the groups overlap), which is why the main text treats the rule as mostly measuring memorisation.

**Caveats.** A single trained model and one sampling run per enzyme (sampling noise alone is up to 1 percentage point elsewhere in this paper). Constitutional recovery, ignoring stereochemistry, at 400 samples is 80.3% (train), 16.5% (val) and 16.8% (test). The reverse of a known reaction appears in the candidate set for 8.1% of test and 6.8% of validation reactions, counted separately from forward hits; for the test set the overlap between the two is in S12.

### S9. Re-ranking Details

All counts are full-reaction recoveries among $n = 2{,}349$ test pairs from the 400-candidate set at $T = 1.0$, whose ceiling is 336 pairs. Frequency features are re-extracted at each budget, so each budget only uses its own samples.

Table 4: Full reactions recovered in the top $k$ by re-ranking policy. The last two rows are the two variants reported in the main paper.

| policy | $k = 1$ | 5 | 10 | 25 | 50 | 100 |
|---|---|---|---|---|---|---|
| sampling order | 2 | 19 | 53 | 121 | 191 | 271 |
| balance only | 8 | 75 | 150 | – | – | – |
| novelty only | 4 | 33 | 83 | – | – | – |
| frequency only | 4 | 66 | 116 | – | – | – |
| novelty + balance | 78 | 198 | 268 | – | – | – |
| balance + frequency (assumption-free) | 5 | 116 | 191 | 297 | 325 | 334 |
| novelty + balance + frequency (novelty-assuming) | 132 | 246 | 291 | 323 | 333 | 334 |

No single feature accounts for the gain. Novelty alone and balance alone both help only a little, and combining them lifts top-1 from 2 pairs to 78. The Wilson 95% interval for the full-reaction top-1 of 5.62% is [4.76, 6.63] and for the top-10 of 12.39% is [11.12, 13.78].

Table 5: Product-set recovery in the top $k$, test $n = 2{,}349$, 400-candidate set at $T = 1.0$: sampling order (before re-ranking) and after re-ranking with the train/validation lookup (novelty-assuming).

| $k$ | before re-ranking | with train/val lookup |
|---|---|---|
| 1 | 19 (0.81%) | 141 (6.00%) |
| 5 | 80 (3.41%) | 291 (12.39%) |
| 10 | 147 (6.26%) | 359 (15.28%) |
| 25 | 279 (11.88%) | 445 (18.94%) |
| 50 | 407 (17.33%) | 507 (21.58%) |

**How the gain scales with the candidate budget.** At every budget, ranking recovers most of what is in the candidate set, and this share falls only slowly as the set grows.

Table 6: Re-ranking against candidate budget $N$. "Ceiling" is the number of pairs whose correct full reaction is present anywhere in the set.

| | | top-1 / top-10 | | | top-10 as |
|---|---|---|---|---|---|
| $N$ | ceiling | sampling order | novelty-assuming | no lookup | % of ceiling |
| 25 | 89 | 2 / 53 | 56 / 89 | 5 / 86 | 100.0 |
| 50 | 141 | 2 / 53 | 81 / 139 | 1 / 125 | 98.6 |
| 100 | 209 | 2 / 53 | 99 / 195 | 3 / 158 | 93.3 |
| 200 | 271 | 2 / 53 | 114 / 244 | 2 / 170 | 90.0 |
| 400 | 336 | 2 / 53 | 132 / 291 | 5 / 183 | 86.6 |

**What balance and the identity filter contribute.** The assumption-free policy also discards candidates whose reactants equal their products. This filters on the candidate alone and is not a ranking signal; its effect is small, and without it top-1, top-5 and top-10 at $N = 400$ are 5, 111 and 183 pairs instead of 5, 116 and 191. The "no lookup" column above also omits it.

Restricting each ranked list to mass-balanced candidates changes nothing at any $k$ for either variant, because all 336 correct full reactions are heavy-atom balanced. Balance does matter for product-set recovery, where the sorted novelty-assuming list recovers 589 pairs at $k = 100$ against 422 for the balanced-only list, since a candidate with the right products and the wrong reactants can still score on the product-set metric.

At $N = 400$, 22.9% of distinct candidates (104,806 of 456,971) but 45.3% of samples are heavy-atom balanced, so the model has learned balance as a soft preference rather than a constraint. Every correct full-reaction candidate is balanced (89 of 89 at $N = 25$, 336 of 336 at $N = 400$), against 27.6% of product-set-correct candidates at $N = 400$. By difficulty class at $N = 400$, the ranked top-1 and top-10 are 129 and 275 for recombination (of 1,275) and 3 and 16 for novel-metabolite pairs (of 1,074).

### S10. Ensemble Union Details

We pooled candidates from three checkpoints with about 133 samples each: the best model (`cbv2_t010`) and two $d_{384}$ models, one initialised with ChemBERTa and one randomly. The union recovers the full reaction for 17.28% of pairs (406) and the product set for 34.01% (799), against 14.30% and 27.84% for 400 samples from the best model alone. Each of the other two models is weaker than the best model at the same sample count, so the gain looks more like the members making partly different errors than like better members.

The union does not help at small $k$, where it matters most. After re-ranking, its top-10 is 280 pairs against 291 for the single model, because the extra correct candidates come with proportionally

Table 7: Recovery from pooled candidates, test $n = 2{,}349$, $T = 1.0$.

| model | samples | product-set | full-reaction |
|---|---|---|---|
| `cbv2_t010` (best) | 133 | 21.24% (499) | 10.39% (244) |
| ChemBERTa-initialised, $d_{384}$ | 133 | 15.58% (366) | 6.39% (150) |
| random-initialised, $d_{384}$ | 133 | 13.28% (312) | 5.41% (127) |
| union of the three | 399 | 34.01% (799) | 17.28% (406) |
| best model alone | 400 | 27.84% (654) | 14.30% (336) |

more wrong ones that look similar to them. At $k = 1, 5, 10, 25, 50$ the single model gives 132, 246, 291, 323, 333 and the union 114, 239, 280, 347, 386, from ceilings of 336 and 406. The union is ahead only beyond $k = 25$, so re-ranking a single model is the better choice when few candidates will be inspected. In sampling order the union's top-10 falls from 53 to 37, because interleaving the models dilutes any single model's ordering. Each member is a single seed and the members differ in both initialisation and architecture, so the diversity cannot be attributed to either.

### S11. Structure Ablation Details

Both models were trained in one job with identical code, hyperparameters (those of `cbv2_t010`), Morgan initialisation and seed, differing only in whether the node features feeding the structural channel were zeroed; the ESM pathway is untouched. Best validation loss was 1.9415 with structure and 1.9442 without.

Table 8: Test recovery within 400 samples ($T = 1.0$) with and without the structural channel, scored on the same pairs.

| metric | with structure | sequence only | only with structure | only without | McNemar $p$ |
|---|---|---|---|---|---|
| full-reaction | 316 (13.45%) | 310 (13.20%) | 124 | 118 | 0.748 |
| product-set | 613 (26.10%) | 633 (26.95%) | 190 | 210 | 0.342 |

Without structure, the identity override collapses to the majority class (validation accuracy 0.003 with structure, 0.997 without), so the zeroed model did lose some of the signal the structural channel carried.

Paired differences are +0.26 points (clustered 95% interval −1.08 to +1.63) for full-reaction and −0.85 points (−2.66 to +0.96) for product-set: opposite in sign, with both intervals spanning zero. The retrained structure model scores 316 against the best checkpoint's 336, which suggests that variation between training runs is about 0.85 points, larger than the effect we are measuring. We used one seed per model. We did not test a smaller conditioning budget or a structure-sensitive subset such as the low-identity bins of S12.

### S12. Additional Checks

**Enzyme-clustered intervals** (4,000 resamples of the 1,277 test enzymes). Clustering by enzyme widens the Wilson intervals only slightly, so repeated enzymes do not explain the reported recovery. Full-reaction at $N = 400$: 14.30%, Wilson [12.95, 15.78], clustered [12.66, 16.05]. Product-set at $N = 400$: 27.93%, Wilson [26.15, 29.78], clustered [25.71, 30.26].

**Recovery by BLAST identity to the nearest training enzyme** (full-reaction, $N = 400$): no hit 0.00% ($n = 3$); $< 30\%$ 13.08% (367); 30–50% 15.00% (920); 50–70% 17.29% (347); 70–90% 13.62% (470); $\geq$90% 10.74% (242). Recovery does not rise with similarity to the training set, and the highest-identity bin is among the lowest.

**Shuffled-enzyme control.** 328 pairs recovered only with the true enzyme and 1 only with the shuffled one; exact McNemar $p = 6 \times 10^{-97}$.

**Rule tiers.** Rows with a matched reaction rule that also occurs in training (1,793) recover at

13.11% [11.62, 14.75]; pairs whose rule is absent from training (8) at 0.00%; pairs with no rule match (548) at 18.43% [15.41, 21.89]. Recovery does not decline on rule-novel pairs, but the unmatched pairs are probably simpler in ways the rule set does not capture, so we draw no conclusion about rule generalisation.

**Reverse-orientation matches** (test, 400 samples): forward 336 pairs, reverse 191, both 146, reverse only 45, forward only 190. Separately, 33 test pairs (1.4%) are exact reverses of a training reaction and 7 more of a validation reaction. Excluding the 33 moves free-running full-reaction recovery from 14.30% to 13.86% (321/2,316) and recombination recovery from 25.02% to 24.48% (304/1,242); 15 of the 336 recoveries fall on those pairs, which are recovered at 45.5% against 13.86% overall. The effect on the headline numbers is small, but recovery on those pairs is elevated.

**Identity and stereo-only reactions.** Five test pairs (0.21%) have reactants equal to products as multisets, of which two are recovered. A further 48 pairs (2.04%) differ only in stereochemistry and would become trivial identities under a constitutional metric, which is why we score with isomeric SMILES.

### S13. Negative Results

We tried four alternatives to the reported design and all did worse, so the main text uses rule-based re-ranking and plain sampling.

**A learned ranker.** A learned ranker over the same re-ranking features did worse than sampling order at top-1 (0 pairs). Positive labels are very rare, 242 of 893,638 candidate entries at $N = 400$, and enzyme-grouped cross-validated average precision was 0.42–0.61. This is the comparison behind the main text's choice of fixed rules over a trained scorer.

**Recombining candidate halves.** Pairing the reactant side of one sample with the product side of another raises the ceiling to 439 pairs against 336, but ranking top-$M$ sides by frequency reaches only 353 pairs at $M = 40$, and its top-1 of 61–67 pairs is well below plain re-ranking's 132. No ranking we tried reached the higher ceiling.

**Mixing temperatures.** Splitting a fixed budget across temperatures is beaten by spending it all at the better one: 200 samples at each of $T = 1.0$ and $T = 1.2$ give 29.33% product-set recovery, against 30.91% for $T = 1.2$ alone.

**Nucleus sampling.** Nucleus sampling gives 16.60% ($p = 0.90$) and 17.84% ($p = 0.95$) product-set recovery within 400 samples, against 27.93% for top-$k = 20$, probably because it narrows an already confident distribution.

Two further limits are practical, not negative results. We did not run budgets above 400 samples, because the sampler runs out of accelerator memory at 1,000 candidates per pair without chunking. So 400 is a ceiling we did not lift, and we have not shown that returns diminish beyond it. A second hyperparameter study did not beat the best configuration (best validation recall 10.11% against 11.05%).

### S14. Recovered Examples

Two sets of recovered full reactions were chosen because they are easy to read, not by rank. Neither is a random sample, so they say nothing about rates, which are in S16. Set 1 is in main paper Figure 4 and set 2 is shown here.

**Set 2: three examples whose product metabolite is absent from training.** A glucosylation of an isoflavone by UDP-glucose, a glucuronidation of an acetylated tryptamine by UDP-glucuronate, and an oxidation of a polyhydroxylated steroid by molecular oxygen. Each is a complete, stereo-specified reaction recovered within 400 samples.

All three examples in set 2 have something in common that may explain why they could be

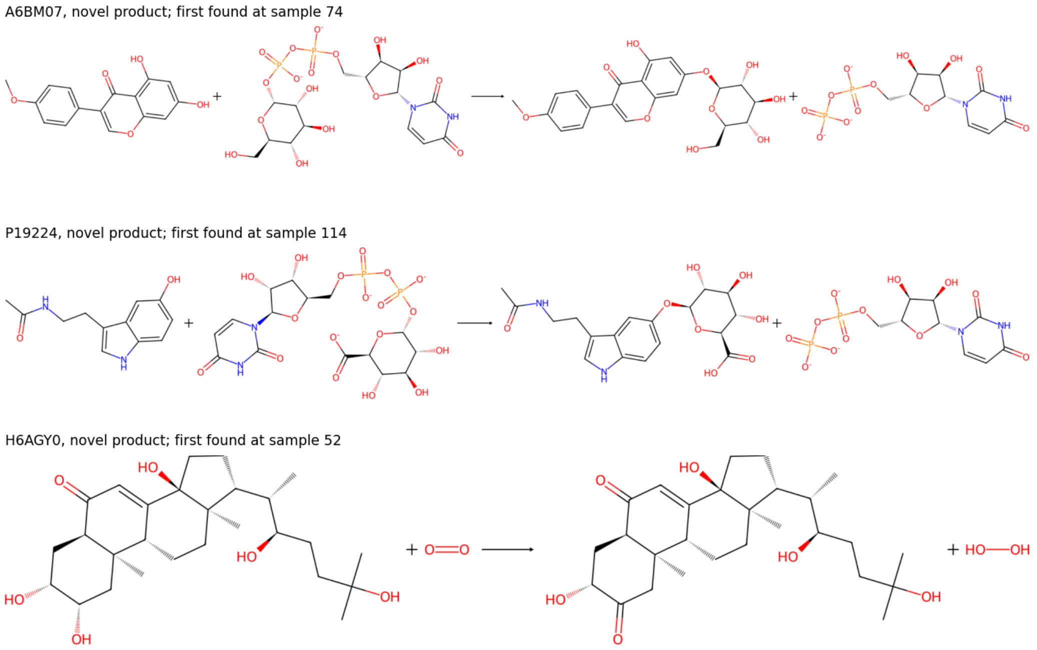


Figure 1: Further recovered full reactions with a product absent from training, with the UniProt entry and the sample at which each first appeared.

recovered. The first two are group transfers whose donor half is a recurring motif in training (UDP-sugar to UDP), so only the acceptor has to be placed, and the acceptors, an isoflavone phenol and a 5-hydroxyindole, are of a kind that glycosyltransferase families commonly accept. The steroid example copies the core unchanged and converts one alcohol to a ketone with $O_2$ reduced to $H_2O_2$, a stoichiometry that recurs across oxidases. These are untested ideas, not established explanations.

### S15. Comparison Tables

These tables support the comparisons in main paper §3.6, which gives the headline values, and add the remaining ranks and the paired differences. The protocols are in S6.

Table 9: Ours vs. a from-scratch RxnPred retrain, paired on the same test pairs.

| **Samples** | **Metric** | **Ours** | **RxnPred retrain** | **Difference** | $p$ |
|---:|---|---:|---:|---:|---:|
| 25 | product-set | 10.34% (243) | 8.09% (190) | +2.26 pp [+0.79, +3.72] | 0.0016 |
| 25 | full-reaction | 4.04% (95) | 3.24% (76) | +0.81 pp [−0.25, +1.88] | 0.14 |
| 50 | product-set | 13.32% (313) | 10.60% (249) | +2.72 pp [+1.16, +4.34] | 0.0006 |
| 50 | full-reaction | 5.96% (140) | 4.73% (111) | +1.23 pp [+0.08, +2.37] | 0.043 |

Table 10: Recall at rank $k$: Horizyn (retrained on EnzymeMap_x) vs. our generator's full-reaction recall after re-ranking, with and without the novelty assumption (main paper §2.3.2). Horizyn ranks a closed library that contains the answer, so the columns are not matched in difficulty.

| $k$ | **Horizyn, retrained** | **Ours, novelty-assuming** | **Ours, assumption-free** |
|---:|---:|---:|---:|
| 1 | 0.77% | 5.62% | 0.21% |
| 25 | 65.30% | 13.75% | 12.64% |
| 100 | 83.95% | 14.22% | 14.22% |

### S16. Recovery Tables

The tables behind main-paper Figure 2 and §3.2–3.3.

Table 11: Recovery within $N$ samples, test set ($n = 2{,}349$), $T = 1.0$, Wilson 95% CI.

| $N$ | **Product-set recovery** | **Full-reaction recovery** |
|---|---|---|
| 1 | 0.81% (19/2,349) [0.52, 1.26] | 0.09% (2/2,349) [0.02, 0.31] |
| 5 | 3.11% (73/2,349) [2.48, 3.89] | 0.68% (16/2,349) [0.42, 1.10] |
| 10 | 5.53% (130/2,349) [4.68, 6.53] | 1.58% (37/2,349) [1.14, 2.16] |
| 25 | 9.37% (220/2,349) [8.25, 10.61] | 3.79% (89/2,349) [3.09, 4.64] |
| 50 | 14.05% (330/2,349) [12.70, 15.51] | 6.00% (141/2,349) [5.11, 7.04] |
| 100 | 19.03% (447/2,349) [17.49, 20.67] | 8.90% (209/2,349) [7.81, 10.12] |
| 200 | 23.54% (553/2,349) [21.87, 25.30] | 11.54% (271/2,349) [10.31, 12.89] |
| 400 | 27.93% (656/2,349) [26.15, 29.78] | 14.30% (336/2,349) [12.95, 15.78] |

Table 12: Recovery within $N$ samples by difficulty class ($T = 1.0$; Wilson 95% CI).

| **Difficulty class ($n$)** | $N$ | **Full-reaction** | **Product-set** |
|---|---|---|---|
| recombination (1,275) | 25 | 6.75% (86) [5.49, 8.26] | 15.14% (193) [13.27, 17.21] |
| recombination (1,275) | 100 | 15.76% (201) [13.87, 17.87] | 29.65% (378) [27.20, 32.21] |
| recombination (1,275) | 400 | 25.02% (319) [22.72, 27.47] | 43.06% (549) [40.37, 45.79] |
| novel_metabolite, all (1,074) | 25 | 0.28% (3) [0.10, 0.82] | 2.51% (27) [1.73, 3.63] |
| novel_metabolite, all (1,074) | 100 | 0.74% (8) [0.38, 1.46] | 6.42% (69) [5.11, 8.05] |
| novel_metabolite, all (1,074) | 400 | 1.58% (17) [0.99, 2.52] | 9.96% (107) [8.31, 11.90] |
| of which novel_product (795) | 25 | 0.38% (3) [0.13, 1.10] | 1.01% (8) [0.51, 1.97] |
| of which novel_product (795) | 100 | 1.01% (8) [0.51, 1.97] | 2.89% (23) [1.94, 4.30] |
| of which novel_product (795) | 400 | 2.14% (17) [1.34, 3.40] | 5.41% (43) [4.04, 7.21] |
| of which novel_reactant_only (279) | 25 | 0.00% (0) [0.00, 1.36] | 6.81% (19) [4.40, 10.39] |
| of which novel_reactant_only (279) | 100 | 0.00% (0) [0.00, 1.36] | 16.49% (46) [12.59, 21.29] |
| of which novel_reactant_only (279) | 400 | 0.00% (0) [0.00, 1.36] | 22.94% (64) [18.39, 28.22] |

### S17. Extended Discussion

The arguments of main paper §4 at greater length.

**What the prior work left open.** Earlier work on RxnPred showed that high reaction-fingerprint similarity does not imply exact or stereo-exact recovery [Balamuhunthan et al., 2026]. What it left unresolved is whether a more geometrically faithful structural encoder would improve exact recovery. That is the question the matched ablation in S11 addresses, and at this scale of supervision we find no measurable effect on exact recovery, though with one seed per model.

**Why both re-ranking variants are reported.** The 14.30% recovered within 400 samples describes the candidate set, not what a user would see. After ranking, the top candidate is the correct full reaction for 5.6% of test pairs and the top ten cover 12.4%, but this assumes that the answer is absent from training, so a ranker can discard every known reaction. That suits a tool that proposes new reactions, but not one that annotates an enzyme whose reaction may already be recorded. For annotation we also report the assumption-free variant, which still gives a 3.6-fold gain at ten candidates. Retrieval models annotate far better than we do, as the Horizyn comparison shows, and a generator can attempt the other task.

**The enzyme-level rule.** Most test enzymes also occur in training, so counting any candidate that matches a known reaction of the enzyme raises apparent recovery from 14.3% to 71.2% (S8). This figure includes memorisation and we do not report it as recovery. It may be closer to what a user would see when applying the model to an enzyme that is already characterised. The numbers we report sit at the low end of this range and are not an estimate of practical usefulness.

### References


Srikrishnan Balamuhunthan, Saketha Nath Jagarlapudi, and Karthik Raman. RxnPred: A generative framework for enzymatic reaction prediction. In *Pacific Symposium on Biocomputing (PSB)*, 2026. Accepted; in press.

Parit Bansal, Anne Morgat, Kristian B. Axelsen, Venkatesh Muthukrishnan, Elisabeth Coudert, Lucila Aimo, Nevila Hyka-Nouspikel, Elisabeth Gasteiger, Arnaud Kerhornou, Teresa B. Neto, et al. Rhea, the reaction knowledgebase in 2022. *Nucleic Acids Research*, 50:D693–D700, 2022. doi: 10.1093/nar/gkab1016.

Bowen Jing, Stephan Eismann, Patricia Suriana, Raphael J. L. Townshend, and Ron Dror. Learning from protein structure with geometric vector perceptrons. In *International Conference on Learning Representations (ICLR)*, 2021.

Radoslav Krivák and David Hoksza. P2Rank: Machine learning based tool for rapid and accurate prediction of ligand binding sites from protein structure. *Journal of Cheminformatics*, 10(1):39, 2018. doi: 10.1186/s13321-018-0285-8.

Greg Landrum. RDKit: Open-source cheminformatics software, version 2025.09.6. https://www.rdkit.org, 2025.

Zeming Lin, Halil Akin, Roshan Rao, Brian Hie, Zhongkai Zhu, Wenting Lu, Nikita Smetanin, Robert Verkuil, Ori Kabeli, Yaniv Shmueli, et al. Evolutionary-scale prediction of atomic-level protein structure with a language model. *Science*, 379:1123–1130, 2023. doi: 10.1126/science.ade2574.

Quinn McNemar. Note on the sampling error of the difference between correlated proportions or percentages. *Psychometrika*, 12:153–157, 1947. doi: 10.1007/BF02295996.

H. L. Morgan. The generation of a unique machine description for chemical structures—a technique developed at chemical abstracts service. *Journal of Chemical Documentation*, 5(2):107–113, 1965. doi: 10.1021/c160017a018.

David Rogers and Mathew Hahn. Extended-connectivity fingerprints. *Journal of Chemical Information and Modeling*, 50(5):742–754, 2010. doi: 10.1021/ci100050t.

Martin Steinegger and Johannes Söding. MMseqs2 enables sensitive protein sequence searching for the analysis of massive data sets. *Nature Biotechnology*, 35:1026–1028, 2017. doi: 10.1038/nbt.3988.

The UniProt Consortium. UniProt: The universal protein knowledgebase in 2023. *Nucleic Acids Research*, 51:D523–D531, 2023. doi: 10.1093/nar/gkac1052.

Mihaly Varadi, Stephen Anyango, Mandar Deshpande, Sreenath Nair, Cindy Natassia, Galabina Yordanova, David Yuan, Oana Stroe, Gemma Wood, Agata Laydon, et al. AlphaFold Protein Structure Database: Massively expanding the structural coverage of protein-sequence space with high-accuracy models. *Nucleic Acids Research*, 50(D1):D439–D444, 2022. doi: 10.1093/nar/gkab1061.
